# Homozygous *CADPS2* mutations cause neurodegenerative disease with Lewy bodies in parrots

**DOI:** 10.1101/2022.03.30.483987

**Authors:** Oswaldo Lorenzo-Betancor, Livio Galosi, Laura Bonfili, Anna Maria Eleuteri, Valentina Cecarini, Ranieri Verin, Fabrizio Dini, Anna-Rita Attili, Sara Berardi, Lucia Biagini, Patrizia Robino, Maria Cristina Stella, Dora Yearout, Michael O. Dorschner, Debby W. Tsuang, Giacomo Rossi, Cyrus P. Zabetian

**Affiliations:** Veterans Affairs Puget Sound Health Care System, Seattle, WA, USA; Department of Neurology, University of Washington School of Medicine, Seattle, WA, USA; School of Biosciences and Veterinary Medicine, University of Camerino, Matelica, Italy; Department of Comparative Biomedicine and Food Science, University of Padova AGRIPOLIS, Legnaro, Italy; Department of Veterinary Sciences, University of Torino, Torino, Italy; Department of Pathology, Center for Precision Diagnostics, University of Washington, Seattle, WA 98195, USA; Department of Psychiatry, University of Washington School of Medicine, Seattle, WA, USA

**Keywords:** *CADPS2*, Lewy body, parkinsonism, parrot, Parkinson’s disease, animal model

## Abstract

**Background:** Several genetic models that recapitulate neurodegenerative features of Parkinson’s disease (PD) exist, which have been largely based on genes discovered in monogenic PD families. However, spontaneous genetic mutations have not been linked to the pathological hallmarks of PD in non-human vertebrates.

**Objective:** To describe the genetic and pathological findings of three yellow crowned parrot (*Amazona ochrocepahala*) siblings with a severe and rapidly progressive neurological phenotype.

**Methods:** The phenotype of the three parrots included severe ataxia, head tilt, and stargazing, while their parents were phenotypically normal. Tests to identify avian viral infections and brain imaging studies were all negative. Due to their inability to survive independently, they were all euthanized at age 3 months and their brains underwent neuropathological examination and proteasome activity assays. Whole genome sequencing (WGS) was performed on the three affected parrots and their parents.

**Results:** The brains of affected parrots exhibited neuronal loss, spongiosis, and Lewy bodies in the neocortex, amygdala, hypothalamus, periaqueductal gray matter, dorsal vagal nucleus, in some cerebellar Purkinje cells, and in the basal ganglia. Proteasome activity was significantly reduced in the affected parrots compared to a control (p<0.05). WGS identified a single homozygous missense mutation (p.V559L) in a highly conserved amino acid residue within the pleckstrin homology (PH) domain of the Calcium Dependent Secretion Activator 2 (*CADPS2*) gene. Previous studies suggest that CADPS2 is expressed at high levels in the substantia nigra where it regulates BDNF release. Thus, disruption of CADPS2 function could impact survival of dopaminergic neurons. Furthermore, CADPS2 expression is in part regulated by two well established PD genes, LRRK2 and SNCA.

**Conclusions:** Our data suggest that a homozygous mutation in the *CADPS2* gene causes a severe neurodegenerative phenotype with Lewy bodies in parrots. Although *CADPS2* variants have not been reported to cause PD in humans, further investigation of the gene in model organisms might provide important insights into the pathophysiology of Lewy body disorders.

## 1. Introduction

The development of genetic animal models that recapitulate Parkinson’s disease (PD) neuropathology has been challenging^1^. Currently, there are several genetic animal models, but none of them completely mimic the clinical findings and associated neuropathological hallmarks of PD^1^.

Several groups have attempted to create α-synuclein transgenic mice with progressive loss of dopamine neurons, including models expressing truncated C-terminus α-synuclein which is more prone to aggregation^2-4^, conditionally expressing either wild type or A53T human α-synuclein^5, 6^, and bigenic crosses with parkin^7^ or DJ-1 knockout mice^8^, but their results have not been promising^4, 9-11^. Of the existing vertebrate models, only the mouse prion promoter (mPrP) A53T α-synuclein transgenic mice model displays the full range of α-synuclein pathology that is observed in human brains, which includes α-synuclein aggregation, fibrils and truncation, α-synuclein phosphorylation and ubiquitination, and progressive age-dependent non-dopaminergic neurodegeneration^12-15^. However, this model does not show progressive degeneration of the dopaminergic system^1^. On the other hand, autosomal recessive knockout models based on *PRKN, PINK1*, or *DJ-1* genes do not show any substantial behavioral or progressive nigrostriatal pathology, nor the typical neuropathological PD hallmark of α-synuclein aggregation^7, 16-23^.

The yellow crowned parrot (*Amazona ochrocephala*) is a species native to tropical Central and South America. Descriptions of neurodegenerative pathologies in parrots and other birds are uncommon, and in most instances the pathology is related to environmental exposures to toxins such as organophosphates and heavy metals that induce axonopathies^24^. Rare cases with central nervous system involvement such as a Lafora disease-like syndrome in cockatiels^25^ and cerebellar degeneration in parrots^26^ and chickens^27^ have been described, but Lewy body pathology has never been reported in birds. Here we present a yellow crowned parrot pedigree with a severe progressive early onset neurological phenotype and widespread Lewy body pathology.

## 2. Material and methods

### 2.1 Animals

Three 3-month-old Yellow Crowned amazons (*Amazona ochrocephala*) were brought to the School of Biosciences and Veterinary Medicine of the University of Camerino in Italy, showing severe neurological symptoms. Subsequently, DNA samples of their parents, two uncles, and their grandparents were acquired using blood collected during routine annual veterinary visits.

### 2.2 Clinical visit and exams

The clinical history of the three affected parrots and their family was collected. From the three affected birds, blood samples were collected at multiple times to perform hematological and biochemical analyses. Given that viral and bacterial infections can cause neurological manifestations in birds, PCR testing for Chlamydophila, Avian Polyomavirus, Avian Bornavirus, Paramyxovirus, Beak and feather disease virus were conducted^28^. Anti-ganglioside antibodies serology was performed to exclude any form of parrot ganglioneuritis^29^. Brain neuroimaging was performed with a veterinary magnetic resonance (MRI) 0.2 Tesla (Esaote S.p.A, Genova, Italy), using T2 (transversal and sagittal), T1 (transversal), FLAIR (dorsal) and STIR (dorsal) sequences.

### 2.3 Pathological analysis

Given that the parrots were unable to feed themselves and the neurological symptoms increased, with eventual death imminent, they were humanitarianly euthanized at the age of 4 months. A complete necropsy was carried out and all organs were fixed in 10% buffered formalin for histological examination. Brain tissue was stained with hematoxylin and eosin, Congo red dye, Luxol fast blue, PAS, and immunohistochemistry (IHC) was performed using an anti-synaptophysin antibody (Agilent Technologies, Inc., Santa Clara, CA, USA), anti-neurofilament antibody (Merck KGaA, Darmstadt, Germany) and an α-synuclein antibody (Santa Cruz Biotechnology, Inc., Dallas, TX, USA). Small portions of brain (1mm^3^) were fixed in 2.5% glutaraldehyde for 24 hours and then in Millonig buffer for electron microscopy. After dehydration the sections were embedded in epoxy resin. Semi-thin toluidine blue 1% stained sections were produced to assess target areas for ultrastructural analysis. Ultra-thin sections (75 nm) were then mounted on copper grids and examined under a Philips EM208S (FEI UK, Cambridge, UK) transmission electron microscope.

To explore the involvement of a CADPS2 mutation in the pathology, IHC was performed with a CADPS2 polyclonal antibody (Invitrogen Corporation, MA, USA), followed by a TUNEL assay. The expression of CADPS2 was analyzed and quantified using ImageJ/Fiji 1.52p software (NIH, USA), as previously reported^30^. As a control for IHC, western blot, and proteasomal analysis, the brain of a healthy parrot of the same species that died without brain lesions, was used.

### 2.4 Western blot and proteasomal analysis

Brains were homogenized in 50 mM Tris buffer, 150 mM KCl, 2 mM EDTA, pH 7.5 (1:5 weight/volume of buffer). Homogenates were immediately centrifuged at 13.000xg for 20 min at 4°C and the supernatant was collected for enzymes activity assays and western blotting. Protein content was determined by the Bradford method^31^ using bovine serum albumin (BSA) as standard. Proteasome peptidase activities in brain homogenates were determined using synthetic fluorogenic peptides: Suc-Leu-Leu-Val-Tyr-AMC was used for chymotrypsin-like (ChT-L) activity, Z-Leu-Ser-Thr-Arg-AMC for trypsin-like (T-L) activity, Z-Leu-Leu-Glu-AMC for peptidyl-glutamyl-peptide hydrolyzing (PGPH) activity^32^. The incubation mixture contained brain homogenates (15 μg total proteins), the proper substrate (5 μM final concentration) and 50 mM Tris–HCl pH 8.0, up to a final volume of 100 μL. Incubation was performed at 37 °C for 60 min and the fluorescence of the hydrolyzed 7-amino-4-methyl-coumarin (AMC) was detected (AMC, λexc = 365 nm, λem = 449 nm; pAB, λexc = 304 nm, λem = 664 nm) on a SpectraMax Gemini XPS microplate reader. The 26S proteasome ChT-L activity was tested including in the reaction mix 10 mM MgCl2, 1 mM dithiothreitol, and 2 mM ATP. Brain homogenates were also analyzed through western blotting assays using anti α-synuclein (C-20) primary antibody (sc-7011, from Santa Cruz Biotechnology, Heidelberg, Germany). The bands were quantified by using a densitometric algorithm. Each Western Blot was scanned (16 bits greyscale) and the obtained digital data were processed through Image J (NIH)^33^ to calculate the background mean value and its standard deviation. The background-free image was then obtained subtracting the background intensity mean value from the original digital data. The integrated densitometric value associated with each band was then calculated as the sum of the density values over all the pixels belonging to the considered band having a density value higher than the background standard deviation. The band densitometric value was then normalized to the relative GAPDH signal intensity. The ratios of band intensities were calculated within the same Western Blot. All the calculations were carried out using the Matlab environment (The MathWorks Inc., MA, USA)^34^.

### 2.5 Whole genome sequencing methods

DNA was extracted from liver of the three affected parrots and from blood of their family members. Whole genome sequencing (WGS) of the three offspring and their parents was performed at the University of Washington with 1 μg of DNA on a HiSeq 2000 Sequencing System (Illumina, San Diego, CA). The WGS data were aligned using a standard BWA pipeline to the Budgerigar (*Melopsittacus undulatus*) reference genome (Budgerigar v6.3)^35^ which contains 25,212 number of scaffolds of an undetermined number of genes. The Budgerigar genome shares more than 99.9% homology with the *Amazona ochrocephala* genome. Annotation was performed with ANNOVAR software^36^ using Ensembl and UCSC gene databases for the Budgerigar v6.3 reference genome. Given the recessive inheritance pattern of the disease (both parents were healthy and related to each other and with their three offspring affected), variant selection was performed as follows: removal of intergenic variants; and subsequent restriction of variants where both parents were heterozygous, and the three offspring were homozygotes. The ortholog variants in humans were annotated according to the gene transcript ENST00000449022.7. The candidate variant identified in the WGS analysis was assessed for co-segregation with disease by Sanger sequencing in all parrots (Figure 1A). Primers were designed using Primer3 v.0.4.0 (https://bioinfo.ut.ee/primer3-0.4.0/) and are available on request.

**Figure 1.**
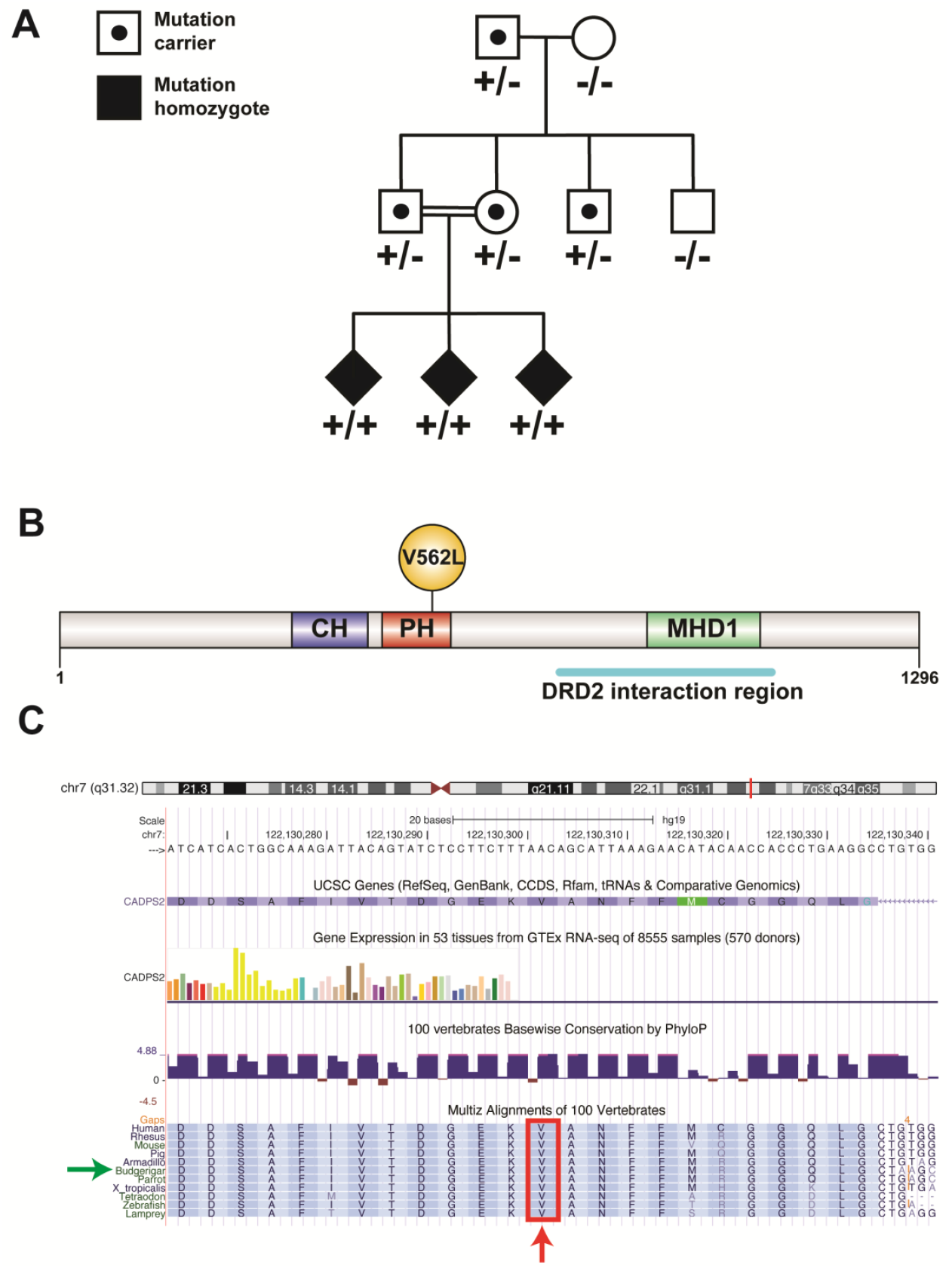
**A. Parrot pedigree**. The grandfather carrying the pathogenic mutation was a wild parrot. The rest of the parrots were born in captivity. Consanguinity of both parents is represented by horizontal double line. **B. Human CADPS2 protein structure**. CH domain = calponin homology domain (family of actin binding domains found in both cytoskeletal proteins and signal transduction proteins); PH domain = the pleckstrin homology domain is a protein domain that occurs in a wide range of proteins involved in intracellular signaling or as constituents of the cytoskeleton; MHD1 domain = the munc13-homology domain 1 may function in a Munc13-like manner to regulate membrane trafficking; DRD2 (dopamine receptor D2) interaction region; **C. Amino acid V562 schematic view that shows its high conservation across species**. Red arrow shows affected valine. Green arrow shows Budgerigar amino acid preservation across species.

## 3. Results

### 3.1 Clinical description

A severe neurological condition was observed in three hand-reared parrot siblings birthed from two different clutches from the same parents, who were siblings (Figure 1A). The grandparents were wild parrots, while the rest of the animals had been born in captivity. The three affected birds hatched from artificially incubated eggs and were hand-fed. They exhibited uncoordinated movements, head tilt, and stargazing (twisted back; see Supplementary video) early in life (2 months). However, the breeder reported that they never exhibited normal behavior, and they were never able to assume a physiological position in the container where they were housed and perform normal movements, compared to parrots of the same age. The early motor signs were stiff neck muscles, and often hyperextension of the limbs, in association with early onset persistent tremor. Tremor was the most noticeable sign observed. It usually began intermittently in one wing and increased considerably when the parrots were under stress or fatigued. The tremor rapidly became bilateral and diffuse, though some degree of asymmetry was still evident. The parrots were unable to perch and they had to be hand-feed despite their age, as they were not able to eat independently. One of the three siblings developed aspiration pneumonia, due to his inability to assume an upright position.

### 3.2 Complementary analyses

Hematological and biochemical findings from blood samples obtained at multiple times were not diagnostic. PCR testing for Avian Polyomavirus, Avian Bornavirus, Paramyxovirus, Beak and feather disease virus and Chlamydophila were all negative. The parrots were also seronegative for anti-ganglioside antibodies. Brain MRI was negative for notable pathology.

On pathological examination, the main microscopical findings observed in the brain included moderate to severe neuronal loss, microspongiosis, and reactive astrogliosis. There were widely distributed variably sized (up to ∼100 µm in diameter), round to elongate, well-defined eosinophilic structures that occasionally contained fine boat-shaped clefts, imparting a crinkled appearance in both gray and white matter. These abundant neuronal and axonal eosinophilic inclusions lacked a distinctive core and halo and resembled Lewy bodies (LBs) and Lewy neurites^37, 38^, which did not stain positively with Congo red, Luxol fast blue, or PAS. Large spherical bodies were occasionally observed in cerebellar Purkinje cells, and multiple small round bodies composed of similar material were noted within pericardial and proventricular ganglia. IHC performed with anti-synaptophysin and neurofilament antibodies failed to stain the eosinophilic bodies. Conversely, strong α-synuclein immunostaining suggested that the round intraneuronal inclusions were consistent with cortical LBs (Fig. 2). These LBs were also present in the neocortex, amygdala, hypothalamus, periaqueductal gray matter, dorsal vagal nucleus, in some cerebellar Purkinje cells, and in the basal ganglia. Some pale bodies and axonal spheroids were present in the same structures. There were no plaques, tangles, or granulovacuolar degeneration in the hippocampal formation. The IHC against α-synuclein confirmed the presence of LBs in the above-mentioned structures. The CADPS2 immunostaining revealed a homogeneously distributed increase of the CADPS2 signal in the affected parrots’ brains when compared to a control (2.5 fold higher than control; Fig. 3D), but LBs did not stain positive for CADPS2 (Fig. 3E).

**Figure 2.**
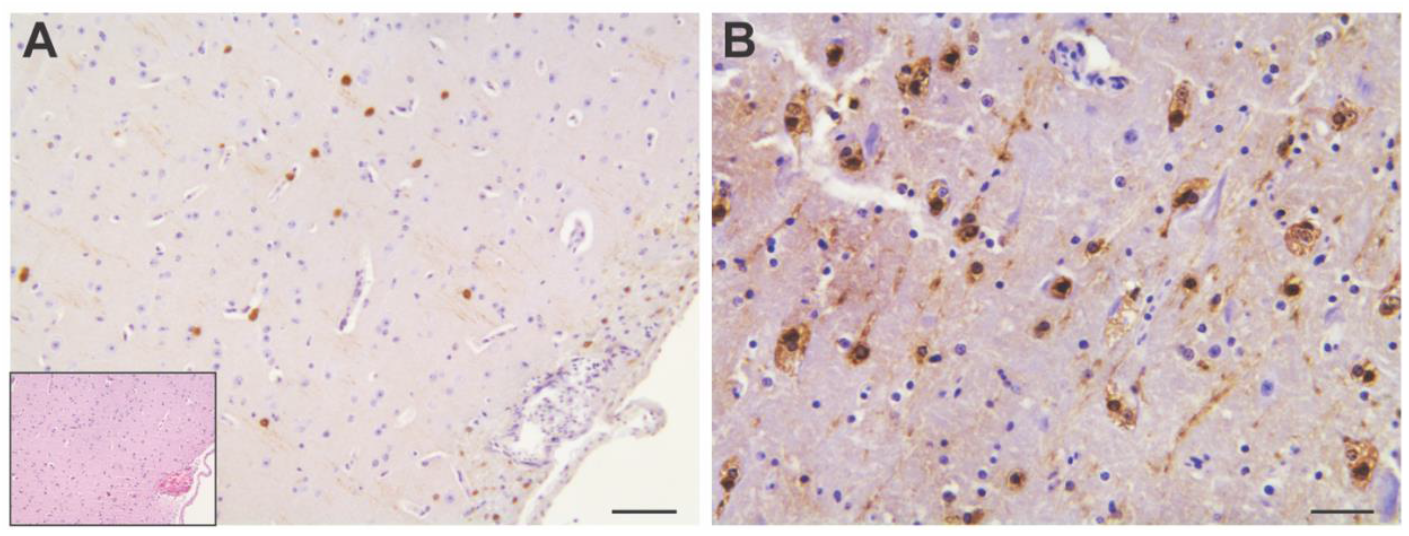
Mutated parrot CNS: midbrain and peri-ventricular area. **A**. In periventricular area some intracytoplasmic inclusions in neurons are immunopositive for α-synuclein (brown staining). In the insert, the same H&E-stained section indicates the sub-meningeal periventricular area in which a scattered inflammatory infiltrate is present. **B**. Higher magnification of the hippocampal area shows a high number of neurons presenting a strongly brown-stained positive cytoplasm for α-synuclein. A vacuolated appearance of the cytoplasm of these neurons is noted. Immunosections were counterstained with Meyer’s hematoxylin. Scale bar A = 250µm, B = 150µm.

**Figure 3.**
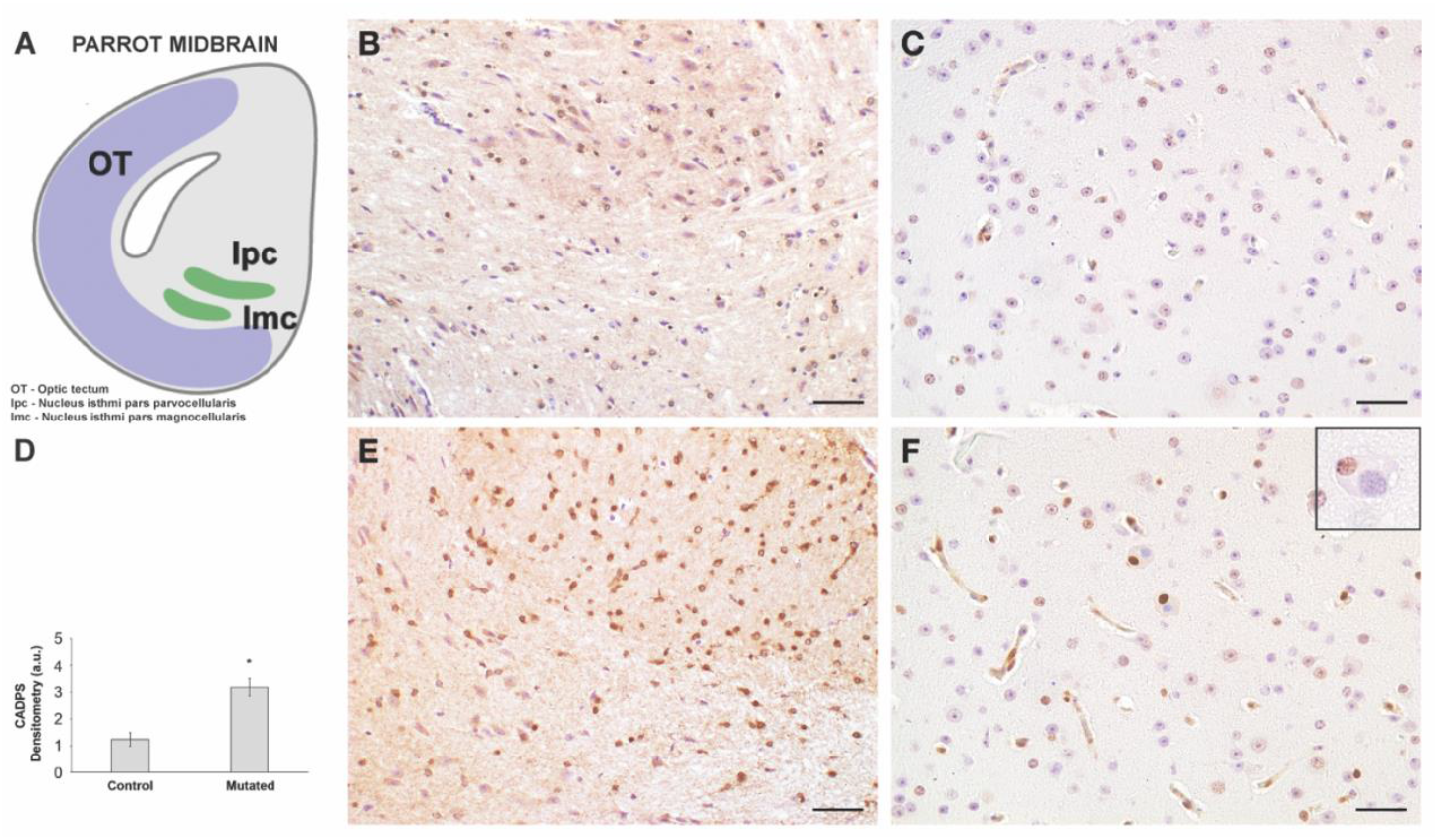
Mutated and control parrot midbrain. **A**. Schematic drawing representing the midbrain region of the birds. **B**. Presence of anti-CADPS2 antibody stain in the periventricular area of the midbrain in a control healthy parrot (**B**) compared to a mutated parrot (**E**). Note the difference of expression and localization, as also quantified in **D**, in the CADPS2 positive neurons (brown stain). **D**. CADPS2 expression analyzed and quantified using ImageJ/Fiji 1.52p software (NIH, USA). The data point marked with an asterisk is statistically significant compared to mutated bird (*p<0,05). **C, F**. Presence of neurons showing TUNEL-positive nucleus in the same parrot’s midbrain region. There is a high concentration of TUNEL positive brown-stained neuronal nuclei in a clear pre-apoptotic state in the section belonging to mutated parrot (**F**), while the TUNEL positive nuclei in the same area of the healthy parrot are few and very lightly stained (**C**). Immunosections were counterstained with Meyer’s hematoxylin. Scale bar B, and E = 250µm; C, and F = 200µm.

In the affected parrots, a conspicuous feature of neurons harboring α-synuclein-positive LBs was somal chromatolytic changes, defined by distension of the cell body, displacement of the nucleus toward the periphery of the soma, and dissolution of the Nissl substance. These neurons also showed nuclear condensation. Subsets of large neurons had this chromatolytic signature, showing TUNEL positive staining (Fig. 3F). Subsets of neurons in brainstem and neocortex were TUNEL positive, indicating cells with double-stranded DNA breaks. In apoptotic neurons, varying degrees of nuclear alterations, ranging from moderate to major chromatin condensation and, in some of these neurons, disappearance of the nucleolus, were observed. Though the nuclear envelope appeared grossly intact it was convoluted. Some of these dying neurons were partially or totally engulfed by glial cells, suggesting an ongoing phagocytic process.

Finally, electron microscopy imaging identified the eosinophilic material constituting LBs as accumulations of short granular electrondense material on a translucid background and organized in short filaments at the periphery (Fig. 4). The presence of an electrondense core, characteristic of LBs in neurons of patients with PD, was not evident in our samples. The material was largely confined to neuronal bodies, and accumulation within the axons was minimal.

**Figure 4.**
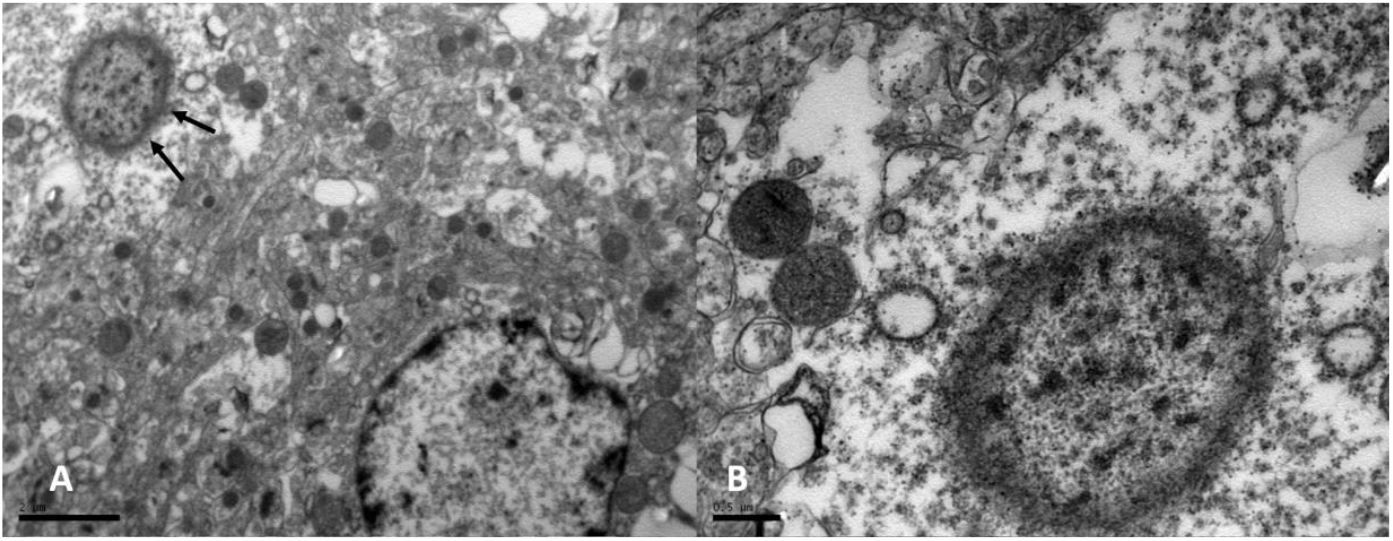
Transmission electron microscopy of a mutated parrot neuron. **A**. Intracytoplasmic LB (black arrows) characterized by accumulations of short granular electrondense material on a translucid background and organized in short filaments at the periphery. **B**. Higher magnification of the same LB. A general loss of mitochondria is observed into neuron, and the two mitochondria near to the LB show an evident loss of cristae and rounded-degenerate appearance. In neurons containing a LB, many mitochondria evidence their spherical pleiomorphism containing poor defined and irregular cristae with finely granular matrix. Some small spherical bodies also appear without cristae. Scale bar A = 2 µm; B = 0.5 µm.

### 3.3 Proteasome and western blot analyses

Significantly reduced ChT-L and T-L proteasomal activities were observed in sick parrots compared to control (p<0.05; Fig. 5A), suggesting a deficit in proteostasis. Moreover, the densitometric analyses obtained from five separate blots detected a significant increase of α-synuclein levels in brain homogenates of each of the three parrots compared to the control (p<0.05; Fig. 5B).

**Figure 5.**
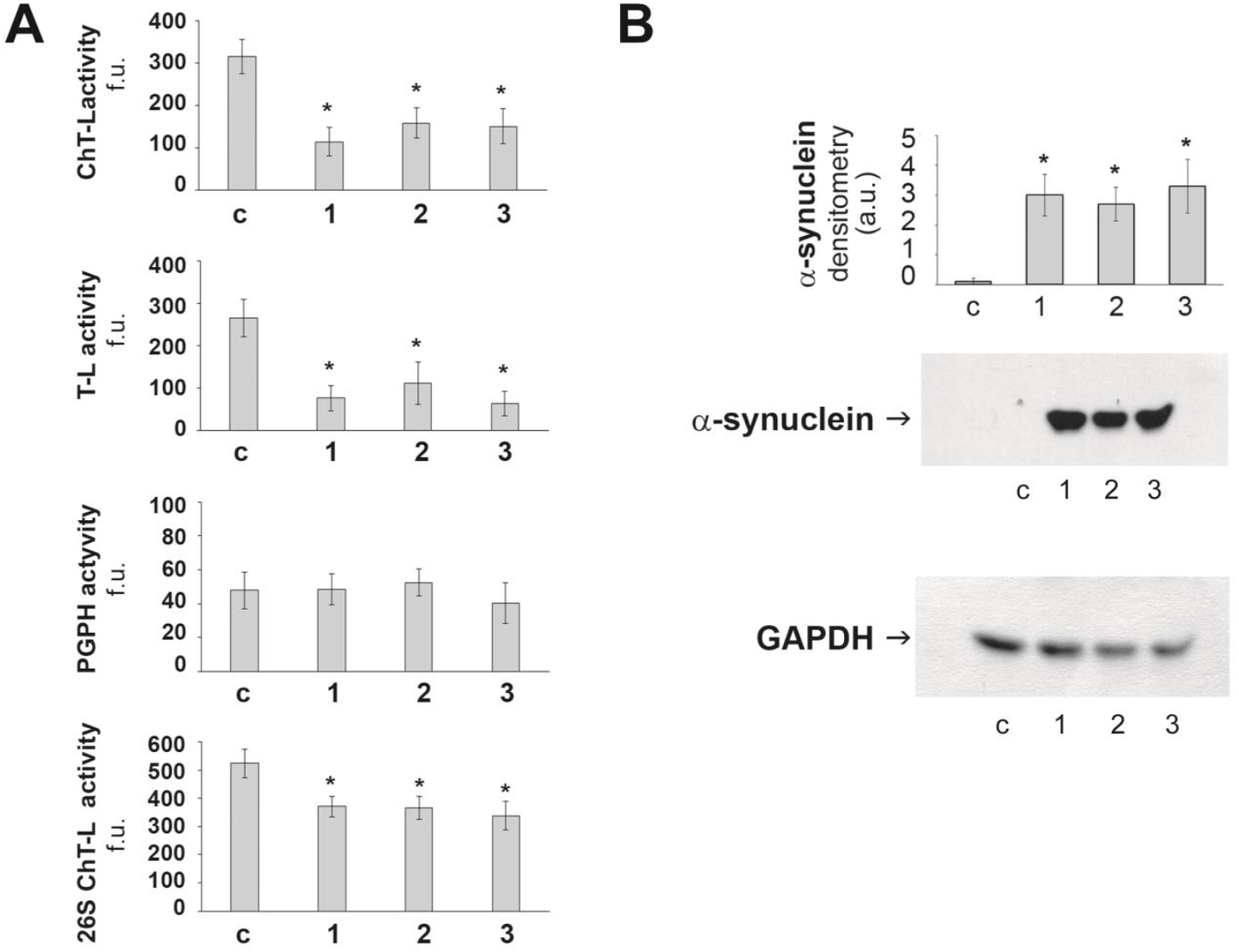
Proteasome activity and α-synuclein levels. **A**. Proteasome activity in control (c) and sick (1,2,3) parrots. 20S proteasome ChT-L, T-L, and PGPH activities and the 26S proteasome ChT-L activity were measured in brain homogenates as described in the Methods section. Results are expressed as fluorescence units (f.u.). Data points marked with an asterisk are statistically significant compared to control bird (*p< 0.05). **B**. Western blot detection of α-synuclein levels in activity in brain homogenates of control (c) and sick (1,2,3) parrots. The densitometric analyses obtained from five separate blots and representative immunoblots are shown. anti-GAPDH antibody was used to check equal protein loading and to normalize target protein. The detection was executed by an ECL western blotting analysis system. Data points marked with an asterisk are statistically significant compared the control bird (*p<0.05).

### 3.4 Whole genome sequencing results

We identified 88 variants genome-wide that matched our filtration criteria (see Supplementary Table 1). Twelve were upstream variants, six were downstream variants, one was located in the 3’ UTR region of a gene, sixty-two were intronic variants, five were synonymous variants and only one was a missense variant that was located in the Calcium Dependent Secretion Activator 2 (*CADPS2*; JH556570.1:3828757C>G; c.1675G>C; p.V559L) gene. The ortholog position and mutation in the human *CADPS2* gene located on chromosome 7 is: g.122130303C>G; c.1684G>C; p.V562L. This amino acid is highly conserved across species, including invertebrates (see Figure 1B and 1C).

## 4. Discussion

In the current paper, we report three parrot siblings with a rapidly progressive neurodegenerative disease caused by a spontaneous homozygous missense mutation (c.1675G>C; p.V559L) in the pleckstrin homology (PH) domain of the *CADPS2* gene. The affected parrots displayed some clinical features of PD including rigidity and tremor though their phenotype was not limited to pure parkinsonism. Their brains displayed widespread neuronal loss and intraneuronal α-synuclein and ubiquitin positive inclusions consistent with LBs, and significant proteasomal dysfunction. The *CADPS2* gene codes for a member of the Calcium Dependent Secretion Activator (CADPS) protein family^39^. This family includes two main genes, *CADPS1*, and *CADPS2*, which are expressed at the highest levels in brain^40-42^. A mouse study showed that CADPS1 regulated catecholamine release from neuroendocrine cells, while CADPS2 regulated the release of two neurotrophins, brain-derived neurotrophic factor (BDNF) and neurotrophin-3 (NT3)^43^. A more recent study of CAPDS2 distribution in mouse brain reported the highest concentrations of CADPS2 immuno-reactivity in the midbrain, cerebellum, and hippocampus. Within the midbrain, peak immunostaining was observed in the cells and mesh-like fiber system of the substantia nigra (SN), ventral tegmental area (VTA), and interpeduncular nucleus (IPN), and overlapped with tyrosine hydroxylase (TH; dopaminergic neuron marker^44^) immunoreactivity in the SN and VTA but not in the IPN. Clear immunoreactivity for CADPS2, but not CADPS1, was substantially localized to TH-positive neurons in mesencephalic-striatal co-cultures^45^. This suggested that CADPS2 is predominantly expressed in dopaminergic neurons of the SN and VTA^45^. In addition, the same study showed that CADPS2 immunostaining overlapped with BDNF-TrkB signaling in the hippocampus^45^.

Taken together, these data suggest that CADPS2 regulates presynaptic BDNF release in dopaminergic neurons, which is particularly relevant to PD and other neurodegenerative diseases. In animal models of PD, BDNF enhances the survival of dopaminergic neurons, improves dopaminergic neurotransmission, and accelerates recovery of motor function^46^. Thus, mutations in CADPS2 might induce neurodegeneration by altering the appropriate release of BDNF.

*CADPS2* is located in the “autism susceptibility locus 1” on chromosome 7q31.32^47^ and genetic variants have been associated with autism spectrum disorders and intellectual disability^48^. Moreover, one isoform of *CADPS2* mRNA lacking exon 3 has been detected in some autistic patients, suggesting that the differential expression pattern of *CADPS2* is involved in neuronal development^49^. Recently, a pooling/bootstrap-based GWAS identified *CADPS2* (rs3757536, *P*=1.54 × 10^−10^) as a modifier of the age of onset in *PSEN1* p.E280A Alzheimer’s disease patients^50^. Two of the most important familial PD related genes are *LRRK2* and *SNCA*^51^. Interestingly, one study performed in cell lines with overexpression of either LRRK2 or α-synuclein showed that CADPS2 expression is differentially regulated by LRRK2 and SNCA^52^. This study showed a significant upregulation (∼2-fold) in both WT and G2019S-LRRK2 expressing cells, when compared to control SHSY5Y cells. Therefore, over-activation of LRRK2, independent of the G2019S mutation, led to increased CADPS2 transcriptional activation suggesting that enhanced LRRK2 cellular function would be sufficient to induce transcriptional dysregulation^52^. The same study evaluated CADPS2 promoter-dependent transcriptional activity in human neuroblastoma SK-N-SH cells overexpressing WT or the PD-causing *SNCA* A30P mutation. In contrast to LRRK2 overexpressing cells, CADPS activity was reduced ∼20% and ∼35% in WT- and A30P cells respectively, when compared to control cells. The effect was again independent from disease causing mutations^52^. Transcriptomic analyses in mice overexpressing human WT α-synuclein have suggested that SNCA may control synaptic vesicle release by downregulating the expression of *Cadps2*^53^, which is critical for constitutive vesicle trafficking and secretion^54^. Thus, CADPS2 expression might be regulated in part by genes that are well-established as playing a role in PD pathogenesis.

In our study we found that parrots harboring the spontaneous homozygous missense mutation (c.1675G>C; p.V559L) had typical hallmarks of neurodegeneration, including intra-neuronal inclusions that resembled LBs, nuclear and cytoplasmic alterations, mitochondrial vacuolization and loss in affected neurons (Figure 4A), and cell death in midbrain. The affected parrots displayed dysmorphic mitochondria (Figure 4B), with some morphological similarities to the shrunken, swollen, or vacuolated mitochondria that have reported in the brains of some PD patients^55^. TUNEL assay identified pro-apoptotic neurons. In brains of affected parrots, neurons with a strong positive TUNEL nucleus were the same ones that showed LB-like intracytoplasmic inclusions (Figure 3F).

Our results suggest that mutations in the *CADPS2* gene cause a severe neurodegenerative phenotype associated with LBs in parrots. Although *CADPS2* variants have not been reported to cause neurodegenerative diseases in humans, further investigation of the gene in animal models might provide important insights into the pathophysiology of LB disorders.

## 5. Acknowledgments

This work was supported by a Merit Review Award from the Department of Veterans Affairs (I01 CX001702) and a Research & Development Seed Grant from the Veterans Affairs Puget Sound Health Care System.

---

**Supplementary table 1.** Mutations identified in the Amazon parrots that met selection criteria.

| Gene ID | Mutation | Human ortholog | Parrots gene name |
| --- | --- | --- | --- |
| **Upstream variants** |  |  |  |
| *ENSMUNG00000007860* | JH556440:1456511 T>G | No orthologs | Not characterized |
| *CCDC3* | JH556467:854099 G>T | *CCDC3* | Coiled-coil domain containing 3 |
| *ENSMUNG00000011465* | JH556513:37413 G>T | No orthologs | Not characterized |
| *ARHGAP29* | JH556574:3047604 G>A | *ARHGAP29* | Rho GTPase activating protein 29 |
| *HES1* | JH556581:130341 T>C | *HES1* | Hes family bHLH transcription factor 1 |
| *SEMA3A* | JH556593:904056 A>G | *SEMA3A* | Semaphorin 3A |
| *MSI2* | JH556627:6313140 AG>A | *MSI2* | Musashi RNA binding protein 2 |
| *MSI2* | JH556627:6313144 T>C | *MSI2* | Musashi RNA binding protein 2 |
| *MSI2* | JH556627:6313148 TCC>T | *MSI2* | Musashi RNA binding protein 2 |
| *MSI2* | JH556627:6313152 TAG>T | *MSI2* | Musashi RNA binding protein 2 |
| *MSI2* | JH556627:6313155 A>T | *MSI2* | Musashi RNA binding protein 2 |
| *MSI2* | JH556627:6313158G C>G | *MSI2* | Musashi RNA binding protein 2 |
| **Downstream variants** |  |  |  |
| *GOLGB1* | JH556230:144106 T>C | *GOLGB1* | Golgin B1 |
| *SELENOF* | JH556574:2880439 A>G | *SELENOF* | Selenoprotein F |
| *KNG1* | JH556579:7170399 G>T | *KNG1* | Kininogen 1 |
| *POMT2* | JH556597:2370780 A>G | *POMT2* | Protein O-mannosyltransferase 2 |
| *CPNE3* | JH556607:8816919 T>A | *CPNE3* | Copine 3 |
| *KCNK12* | JH556611:3308002 A>G | *KCNK12* | Potassium two pore domain channel subfamily K member 12 |
| **3' UTR region** |  |  |  |
| *ENSMUNG00000009678* | JH556472:111791T>C | No orthologs | Dual specificity testis-specific protein kinase 1-like |
| **Intronic variants** |  |  |  |
| *HPS3* | JH556131:245809 T>C | *HPS3* | HPS3 biogenesis of lysosomal organelles complex 2 subunit 1 |
| *RNF141* | JH556236:326470 T>G | *RNF141* | Ring finger protein 141 |
| *EFTUD2* | JH556240:20215 T>C | *EFTUD2* | Elongation factor Tu GTP binding domain containing 2 |
| *RNF123* | JH556294:363891 G>T | *RNF123* | Ring finger protein 123 |
| *CACNA2D2* | JH556294:1600415 T>G | *CACNA2D2* | Ca<sup>2+</sup> voltage-gated channel auxiliary subunit alpha 2 delta 2 |
| *EIF2B5* | JH556422:124552 T>C | *EIF2B5* | Eukaryotic translation initiation factor 2B subunit epsilon |
| *SFXN1* | JH556443:1749967 C>T | *SFXN1* | Sideroflexin 1 |
| *PPM1E* | JH556455:912740 T>C | *PPM1E* | Protein phosphatase, Mg<sup>2+</sup>/Mn<sup>2+</sup> dependent 1E |
| *PPP2R3A* | JH556459:721355 G>C | *PPP2R3A* | Protein phosphatase 2 regulatory subunit B, alpha |
| *EPHB1* | JH556459:1214019 G>T | *EPHB1* | EPH receptor B1 |
| *RYK* | JH556459:2115989 A>G | *RYK* | Receptor like tyrosine kinase |
| *CCDC3* | JH556467:811114 G>A | *CCDC3* | Coiled-coil domain containing 3 |
| *OPTN* | JH556467:878281 G>A | *OPTN* | Optineurin |
| *ENSMUNG00000015055* | JH556467:1062317 G>C | No orthologs | pre-mRNA processing factor 18 |
| *DMD* * | JH556496:339158 T>C* | *DMD* | Dystrophin |
| *MGMT* | JH556537:1071186 G>A | *MGMT* | O-6-methylguanine-DNA methyltransferase |
| *LTBP1* | JH556557:3812706 C>T | *LTBP1* | Latent transforming growth factor beta binding protein 1 |
| *BRINP1* | JH556563:892106 T>G | *BRINP1* | BMP/retinoic acid inducible neural specific 1 |
| *FANCD2* | JH556566:2038768 A>G | *FANCD2* | FA complementation group D2 |
| *CNST* | JH556569:6571419 A>G | *CNST* | Consortin, connexin sorting protein |
| *MET* | JH556570:1320112 T>C | *MET* | MET proto-oncogene, receptor tyrosine kinase |
| *CPED1* | JH556570:3154117 T>A | *CPED1* | Cadherin like and PC-esterase domain containing 1 |
| *CADPS2* | JH556570:3827987 G>A | *CADPS2* | Calcium dependent secretion activator 2 |
| *GRM8* | JH556570:5483008 T>C | *GRM8* | Glutamate metabotropic receptor 8 |
| *GRM8* | JH556570:5505143 C>T | *GRM8* | Glutamate metabotropic receptor 8 |
| *GRM8* | JH556570:5505154 G>A | *GRM8* | Glutamate metabotropic receptor 8 |
| *GRM8* | JH556570:5711450 G>T | *GRM8* | Glutamate metabotropic receptor 8 |
| *UTS2B* | JH556574:1593897 T>C | *UTS2B* | Urotensin 2B |
| *GOLIM4* | JH556579:1719630 T>C | *GOLIM4* | Golgi integral membrane protein 4 |
| *NAALADL2* | JH556579:4533435 A>G | *NAALADL2* | N-acetylated alpha-linked acidic dipeptidase like 2 |
| *VOPPI* | JH556580:1511178 C>T | *VOPPI* | WW domain binding protein |
| *ADAMTS2* | JH556582:2582462 G>A | *ADAMTS2* | ADAM metalloproteinase with thrombospondin type 1 motif 2 |
| *PLCB1* | JH556583:7325538 A>T | *PLCB1* | Phospholipase C beta 1 |
| *YDJC* | JH556586:4414982 A>G | *YDJC* | YdjC chitoooligosaccharide deacetylase homolog |
| *CAMSAP2* | JH556589:5850269 T>C | *CAMSAP2* | Calmodulin regulated spectrin assoc protein fam member 2 |
| *SEMA3A* | JH556593:981396 A>G | *SEMA3A* | Semaphorin 3A |
| *NAPEPLD* | JH556593:4793469 T>C | *NAPEPLD* | N-acyl phosphatidylethanolamine phospholipase D |
| *PUS7* | JH556593:5909299 C>T | *PUS7* | Pseudouridine synthase 7 |
| *PRKAR2B* | JH556593:6494204 A>G | *PRKAR2B* | Protein kinase cAMP-dependent type II regulatory subunit beta |
| *CD151* | JH556599:2918486 C>T | *CD151* | CD151 molecule (Raph blood group) |
| *NOVA1* | JH556599:9765500 C>G | *NOVA1* | NOVA alternative splicing regulator 1 |
| *EGLN3* | JH556599:12934752 A>G | *EGLN3* | Egl-9 family hypoxia inducible factor 3 |
| *ENSMUNG00000015432* | JH556600:9258645 A>G | No orthologs | Forkhead box P1 |
| *SUMF1* | JH556600:12184762 T>C | *SUMF1* | Sulfatase modifying factor 1 |
| *ENSMUNG00000014258* | JH556605:2875840 C>T | No orthologs | Guanylate cyclase soluble subunit beta-2-like |
| *LARGE1* | JH556605:13276382 G>A | *LARGE1* | LARGE xylosyl- and glucuronyltransferase 1 |
| *RDH10* | JH556607:3245659 T>G | *RDH10* | Retinol dehydrogenase 10 |
| *SDC2* | JH556607:12770380 G>A | *SDC2* | Syndecan 2 |
| *SLC44A5* | JH556608:18578303 T>C | *SLC44A5* | Solute carrier family 44 member 5 |
| *CDC42BPA* | JH556611:4757071 C>T | *CDC42BPA* | CDC42 binding protein kinase alpha |
| *TGFB2* | JH556611:7687237 A>G | *TGFB2* | Transforming growth factor beta 2 |
| *ENSMUNG00000000832* | JH556614:7806740 T>C | No orthologs | Not characterized |
| *SCUBE2* | JH556615:23817043 G>A | *SCUBE2* | Signal peptide, CUB domain and EGF like domain containing 2 |
| *GTF2H1* | JH556615:27192731 A>G | *GTF2H1* | General transcription factor IIH subunit 1 |
| *ENSMUNG00000013397* | JH556617:30408533 G>C | No orthologs | Unconventional myosin-X-like |
| *PDLIM1* | JH556617:31042012 C>T | *PDLIM1* | PDZ and LIM domain 1 |
| *TGFBR3* | JH556617:34306232 C>T | *TGFBR3* | Transforming growth factor beta receptor 3 |
| *SPSB4* | JH556617:35184609 A>G | *SPSB4* | SplA/ryanodine receptor domain and SOCS box containing 4 |
| *ENSMUNG00000014151* | JH556617:37077077 A>G | No orthologs | Glypican-5-like |
| *LIMS2* | JH556617:39010271 T>G | *LIMS2* | LIM zinc finger domain containing 2 |
| *DPYD* | JH556617:39665970 A>C | *DPYD* | Dihydropyrimidine dehydrogenase |
| *VTI1A* | JH556628:6394349 C>T | *VTI1A* | Vesicle transport through interaction with t-SNAREs 1A |
| **Synonymous variants** |  |  |  |
| *SPEN* | JH556559:383406 G>T | *SPEN* | Spen family transcriptional repressor |
| *POT1* | JH556570:4837014 T>C | *POT1* | Protection of telomeres 1 |
| *ENSMUNG00000011074* | JH556581:464702 C>T | No orthologs | Protein phosphatase 1 regulatory inhibitor subunit 2 |
| *MYOM1* | JH556616:28872447 G>A | *MYOM1* | Myomesin 1 |
| *ENSMUNG00000008400* | JH556617:7477768 A>T | Not orthologs | Tet methylcytosine dioxygenase 1 |
| **Missense variant** |  |  |  |
| ***CADPS2*** | **JH556570:3828757 C>G** | ***CADPS2*** | **Calcium dependent secretion activator 2** |

\* The ortholog position in human *DMD* gene is intronic, too.

